# Prader-Willi locus *Snord116* RNA processing requires an active endogenous allele and neuron-specific splicing by *Rbfox3/NeuN*

**DOI:** 10.1101/305557

**Authors:** Rochelle L. Coulson, Weston T. Powell, Dag H. Yasui, Gayathri Dileep, James Resnick, Janine M. LaSalle

## Abstract

Prader-Willi syndrome (PWS), an imprinted neurodevelopmental disorder characterized by metabolic, sleep, and neuropsychiatric features, is caused by the loss of paternal *SNORD116*, containing only noncoding RNAs. The primary *SNORD116* transcript is processed into small nucleolar RNAs (snoRNAs), which localize to nucleoli, and their spliced host gene *116HG*, which is retained at its site of transcription. While functional complementation of the *SNORD116* noncoding RNAs is a desirable goal for treating PWS, the mechanistic requirements of *SNORD116* RNA processing are poorly understood. Here we developed and tested a novel transgenic mouse which ubiquitously expresses *Snord116* on both a wild-type and *Snord116* paternal deletion *(Snord116^+/−^)* background. Interestingly, while the *Snord116* transgene was ubiquitously expressed in multiple tissues, splicing of the transgene and production of snoRNAs was limited to brain tissues. Knockdown of *Rbfox3*, encoding neuron-specific splicing factor NeuN, in Snord116^+/−^-derived neurons reduced splicing of the transgene in neurons. RNA fluorescent *in situ* hybridization for *116HG* revealed a single significantly larger signal in transgenic mice, demonstrating colocalization of transgenic and endogenous *116HG* RNAs. Similarly, significantly increased snoRNA levels were detected in transgenic neuronal nucleoli, indicating that transgenic *Snord116* snoRNAs were effectively processed and localized. In contrast, neither transgenic *116HG* nor snoRNAs were detectable in either non-neuronal tissues or *Snord116^+/−^* neurons. Together, these results demonstrate that exogenous expression and neuron-specific splicing of the *Snord116* locus are insufficient to rescue the genetic deficiency of *Snord116* paternal deletion. Elucidating the mechanisms regulating *Snord116* processing and localization are essential to develop effective gene replacement therapies for PWS.

## Introduction

Prader-Willi syndrome (PWS) is a neurodevelopmental disorder characterized by a broad range of symptoms including hypotonia and failure to thrive in infancy followed by the onset of hyperphagia, intellectual impairment, obsessive-compulsive tendencies, and sleep abnormalities including shorter sleep duration and daytime sleepiness (1). PWS is caused by paternal deficiency of the maternally imprinted 15q11-q13 locus, which encodes a neuron-specific ~1Mb transcript (Fig. 1). Expression of this locus originates at the PWS imprinting control region (IC) at the 5’ end of *SNRPN*, extends through small nucleolar RNA (snoRNA) clusters *Snord116* and *Snord115*, and terminates antisense to the maternally expressed *Ube3a (Ube3a-ats)* (2–5). Analyses of PWS patients have revealed that small deletions of the *Snord116* cluster of snoRNAs are sufficient to cause PWS (6–9). *Snord116* is processed to form two noncoding RNAs: *Snord116* snoRNAs and the *Snord116* host gene *(116HG). Snord116* snoRNAs are intronically embedded within the noncoding *Snord116* primary transcript, and although they currently have no known target sequence, they localize to the nucleolus in mature neurons (10–12), detectable by RNA fluorescence *in situ* hybridization (FISH) (Fig. 1). It has also been proposed that these orphan snoRNAs are further processed, generating short fragments with non-canonical functions. Generation of these processed snoRNAs (psnoRNAs) is not well understood, however it has been suggested that they may arise from the same host intron and depend on stability and protein interactions (13). The nucleolar accumulation of *Snord116* snoRNAs coincides with increased transcription of the locus, an increase in nucleolar size during early postnatal development, and chromatin decondensation of the paternally expressed *Snrpn-Ube3a* allele, detected by DNA FISH (10).

**Figure 1.**
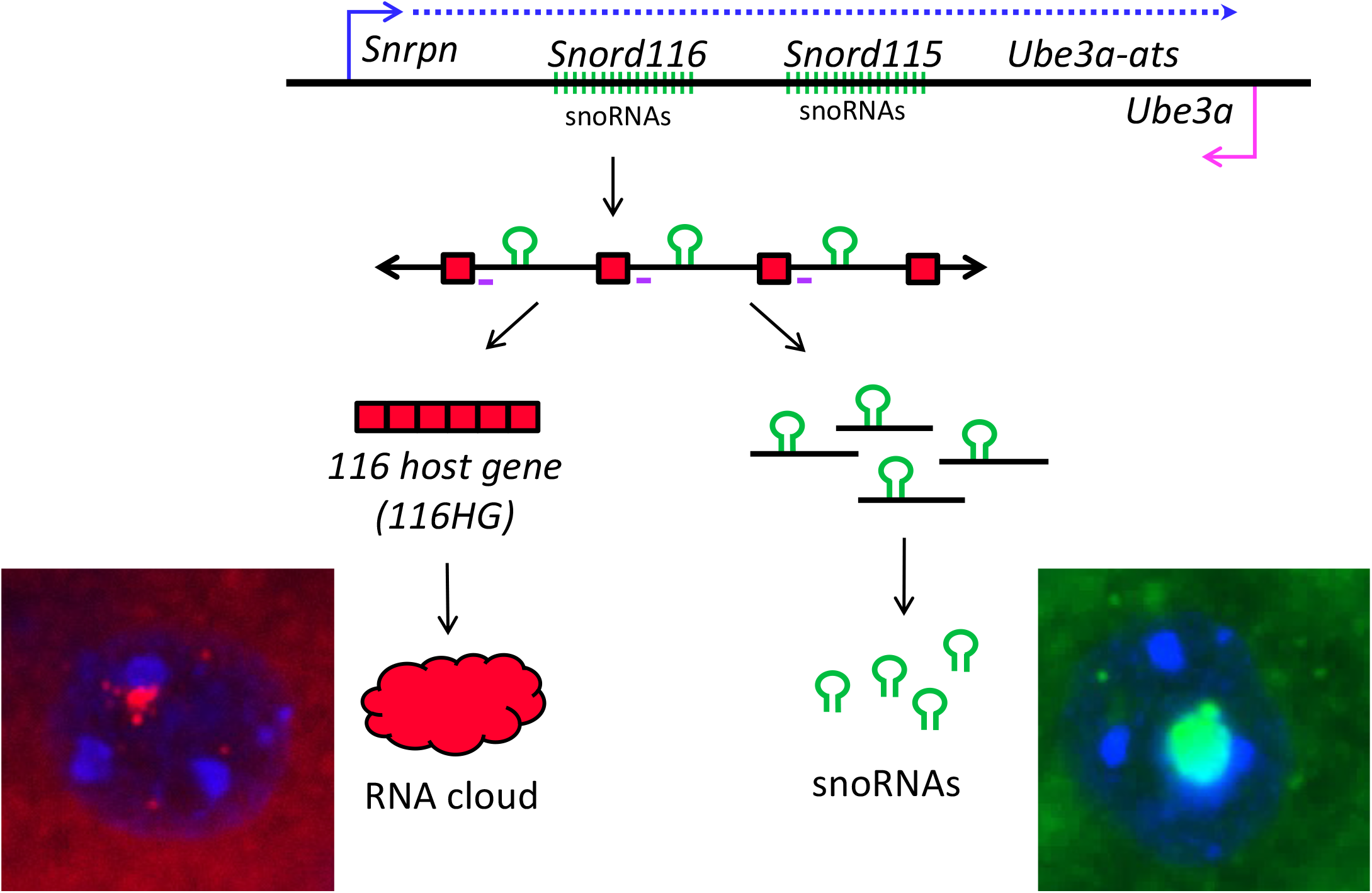
The PWS locus encodes a long ˜1 Mb transcript containing the Snordll6 and Snordll5 snoRNA clusters. Processing of this primary transcript produces snoRNAs from the introns of *Snord116* and the *116HG* from the spliced exons of *Snord116.* Localization of the *116HG* and *Snord116* snoRNAs are shown by RNA FISH. Location of Rbfox3 binding motifs indicated on primary transcript in purple.

*116HG* is a long noncoding RNA (lncRNA) generated from the spliced exons of the *Snord116* primary transcript. *116HG* is retained within the nucleus and localizes to its paternally decondensed site of transcription, forming an “RNA cloud”, which associates with the *Snord116* locus as well as other genes with epigenetic, metabolic, and circadian functions, regulating their expression in a time-of-day dependent manner (11). The *116HG* cloud is significantly larger during light hours, corresponding to the downregulated expression of its gene targets during sleep (14). Co-localized nuclear accumulation of the unspliced transcript and the spliced *116HG* indicates that splicing occurs locally at the site of paternal expression (11). Although splicing is required for *116HG* formation and *Snord116* snoRNA biogenesis, the mechanism of this process and its specificity for neurons is not well understood.

In an effort to determine phenotypes associated with genes within the PWS region, mouse models carrying deletions of nearly every paternally expressed gene in the human 15q11-q13 locus, either individually or as part of a large deletion, have been created and characterized (15). Although some models exhibit a subset of PWS-like phenotypes, issues of lethality complicate phenotypic analysis of adult mice in some models, and no current mouse model of PWS exhibits consistency in the late onset obesity characteristic of PWS (15–17). Currently, two mouse models *(PWScr^m+/p-^* and *Snord116^+/−^)* carry deletions of the minimal *Snord116* PWS critical region, and display postnatal growth deficiency characteristic of the early failure to thrive phenotype exhibited by PWS patients as well as some hyperphagic behavior (18,19). Activation of maternal *Snord116* expression rescued the growth retardation and postnatal lethality phenotypes of the *PWScrm^+/p-^* small deletion model, supporting *Snord116* as the critical PWS region (20). Although the processed snoRNAs have previously been the main focus of studies of the *Snord116* locus, transgenic expression of a single copy of *Snord116* snoRNA failed to rescue the phenotype of a *Snrpn-Ube3a* deletion mouse model, suggesting either that the *Snord116* functional unit is not restricted solely to the snoRNAs or that multiple copies are required (19,21). Re-introduction of multiple copies of *Snord116* snoRNAs expressed from the introns of another host gene failed to rescue the growth retardation phenotype of *PWScr^m+/p-^* mice, highlighting the functional significance of the *116HG* (20). Importantly, the regulation of circadian and metabolic gene expression by *116HG* leads to dysregulated energy expenditure in mice deficient for paternal *Snord116*, suggesting that the lncRNA *116HG* may play role in the pathogenesis of PWS (14). Finally, the introns of the *Snord116* primary transcript may play a role in the regulation of *Ube3a-ats* progression by regulating chromatin compaction through the formation of R-loops (22). These studies have illustrated the genetic complexity of the *Snord116* locus and the potential functional capacity of multiple elements within this long noncoding RNA.

We designed a novel transgenic *Snord116* mouse to characterize the processing and formation of these *Snord116* RNA products, and define the mechanism regulating the brain specificity of these processes. By driving expression of *Snord116* with a ubiquitous promoter, we investigated the requirements for *Snord116* processing in multiple tissues and the potential of our transgene to compensate for the molecular deficits observed in *Snord116^+/−^* mice. We show that *Snord116* expression is not sufficient for the production of snoRNAs or the *116HG*, and that the formation of these products is blocked at the level of tissue-specific splicing. Our data demonstrate a requirement for Rbfox3 in the neuron-specific splicing of *Snord116*, and an active paternal *Snord116* allele in the localization of its processed RNA components, providing a better understanding of the requirements of *Snord116* function in the development of potential future therapies for PWS.

## Results

### Transgenic *Snord116* integrates into the genome and is expressed in many tissues

To reintroduce *Snord116* in a highly expressed, ubiquitous manner to paternally Snord116-deficient mice, we engineered a novel “complete” *Snord116* transgene containing three complete *Snord116* repeats under the control of a CMV promoter (Fig. 2a). Each unit of the *Snord116* repeat contained an intronically encoded snoRNA flanked by *116HG* exons, as organized in the genomic DNA. This construct was randomly inserted into the genome by pronuclear injection of fertilized oocytes to create a transgenic mouse carrying 9 copies of the construct, representing 27 copies of the *Snord116* repeat unit (Fig. 2b). Mice were tested for transgene insertion by PCR using primers specific to the transgene and inverse PCR was performed to identify the genomic location of the transgene integration at 7qE3, approximately 47 Mb from the endogenous *Snord116* locus (Fig. 2c). This falls outside of the 15q11-q13 imprinted locus and is therefore not expected to be co-regulated with endogenous *Snord116.*

**Figure 2.**
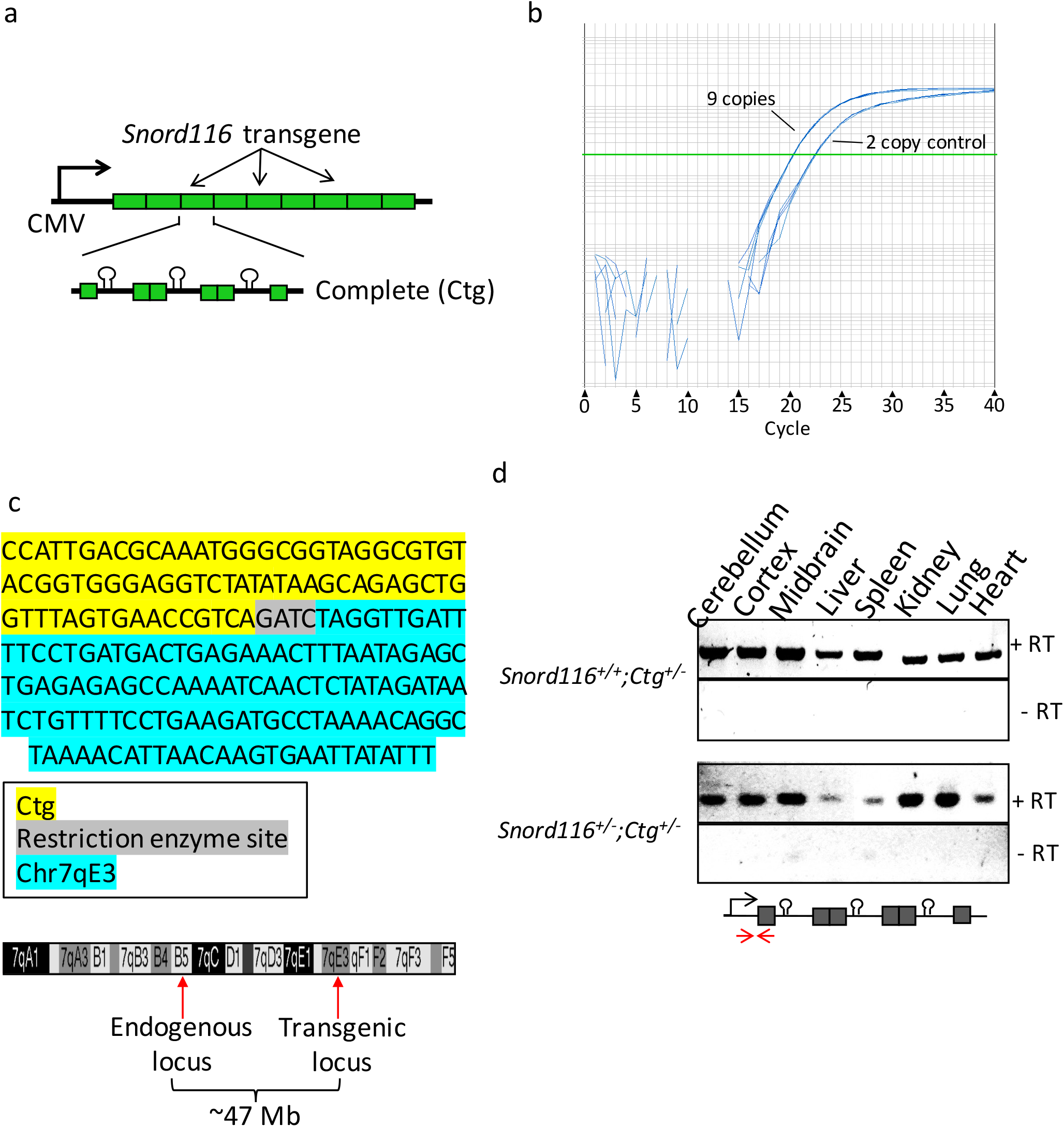
The *Snord116* transgene contains 27 copies of the *Snord116* repeat unit and is ubiquitously expressed. **(a)** Schematic of the *Snord116* transgene construct design containing 9 copies of the 3 copy *Snord116* repeat unit, **(b)** Copy number analysis indicates insertion of the *Snord116* transgene nine times in *Snord116^+/+^;Ctg^+/−^* transgenic mice, **(c)** The location of *Snord116* transgene insertion was identified at chromosome 7qE3 by inverse PCR. Map of chromosome 7 indicating the locations of the endogenous and transgenic loci ~47 Mb apart, **(d)** The *Snord116* transgene shows ubiquitous expression in all tissues tested using transgene-specific primers on both WT and *Snord116^+/^* backgrounds.

Transgenic *Snord116* insertion overlaps with the poorly characterized gene *Gm1966*, which has been reported as both an unprocessed pseudogene as well as a protein coding gene in different databases.

The *Snord116^+/−^* PWS mouse model carries a ~150 kb deletion of *Snord116*, representing the smallest region of overlap for human PWS deletions (6,7,9,19). *Snord116+^-^* mice recapitulate the neonatal failure to thrive exhibited by PWS patients and exhibit altered energy expenditure (14,19). We crossed *Snord116^+/+^;Ctg^+/−^* transgenic females to *Snord116^+-^* males to produce mice deficient for endogenous paternal *Snord116* but carrying the transgene *(Snord116^+/−^;Ctg^+/−^)* and littermate controls *(Snord116^+/+^;Ctg^+/−^).* To ensure that the *Snord116* transgene was not transcriptionally silenced, we tested the expression of the transgene in both WT *(Snord116^+/+^;Ctg^+/−^)* and PWS *(Snord116^+/−^;Ctg^+/−^)* backgrounds, confirming expression in several tissues, including those in which *Snord116* is not endogenously expressed (12) (Fig. 2d).

### Transgenic *Snord116* snoRNAs localize with endogenous snoRNAs specifically in neuronal nucleoli in wild-type but not *Snord116^+/−^* mice

*Snord116* expression is restricted to neurons in mice, however we sought to determine if transgenic *Snord116* was sufficient to recruit the required processing factors for the generation of functional snoRNAs. The ubiquitous expression pattern of the *Snord116* transgene allowed us to examine the processing and localization of *Snord116* snoRNAs outside of the neuronal lineage. We examined *Snord116* snoRNA localization by RNA FISH in coronal brain sections using probes that detect both the endogenous and transgenic *Snord116* RNAs (Fig. 3a and Figs. S1 and S2). Despite ubiquitous expression of the *Snord116* transcript in multiple tissues, detection of snoRNA nucleolar localization was limited to neurons in the brain, as it was not observed in non-neuronal nuclei. *Snord116* snoRNAs were clearly detected in WT nucleoli of Purkinje neurons in the cerebellum and localized to the single large nucleolus. As expected, snoRNAs were not observed in *Snord116+^-^* neurons due to the lack of paternal *Snord116* expression. In *Snord116^+/+^;Ctg^+/−^* mice, the intensity of the nucleolar snoRNA RNA FISH signal was significantly greater than in WT nucleoli (*Snord116^+/+^;Ctg^−/−^*) (Fig. 3b), indicating that transgenic *Snord116* potentially contributes to increasing the snoRNA population. SnoRNAs were also detected in the cortex of WT and *Snord116^+/+^;Ctg^+/−^* mice. However, in the absence of endogenous *Snord116 (Snord116^+/−^;Ctg^+/−^)*, expression of the transgene was not sufficient to produce detectable snoRNA accumulation within neuronal nucleoli in any brain region or in nonneuronal tissues (Figs. S1 and S2). These results indicate that although transgenic *Snord116* is able to contribute to endogenous snoRNAs, transcription of the primary transcript is not sufficient for the processing or localization of these RNAs in non-neuronal and *Snord116*-deficient neuronal tissues.

**Figure 3.**
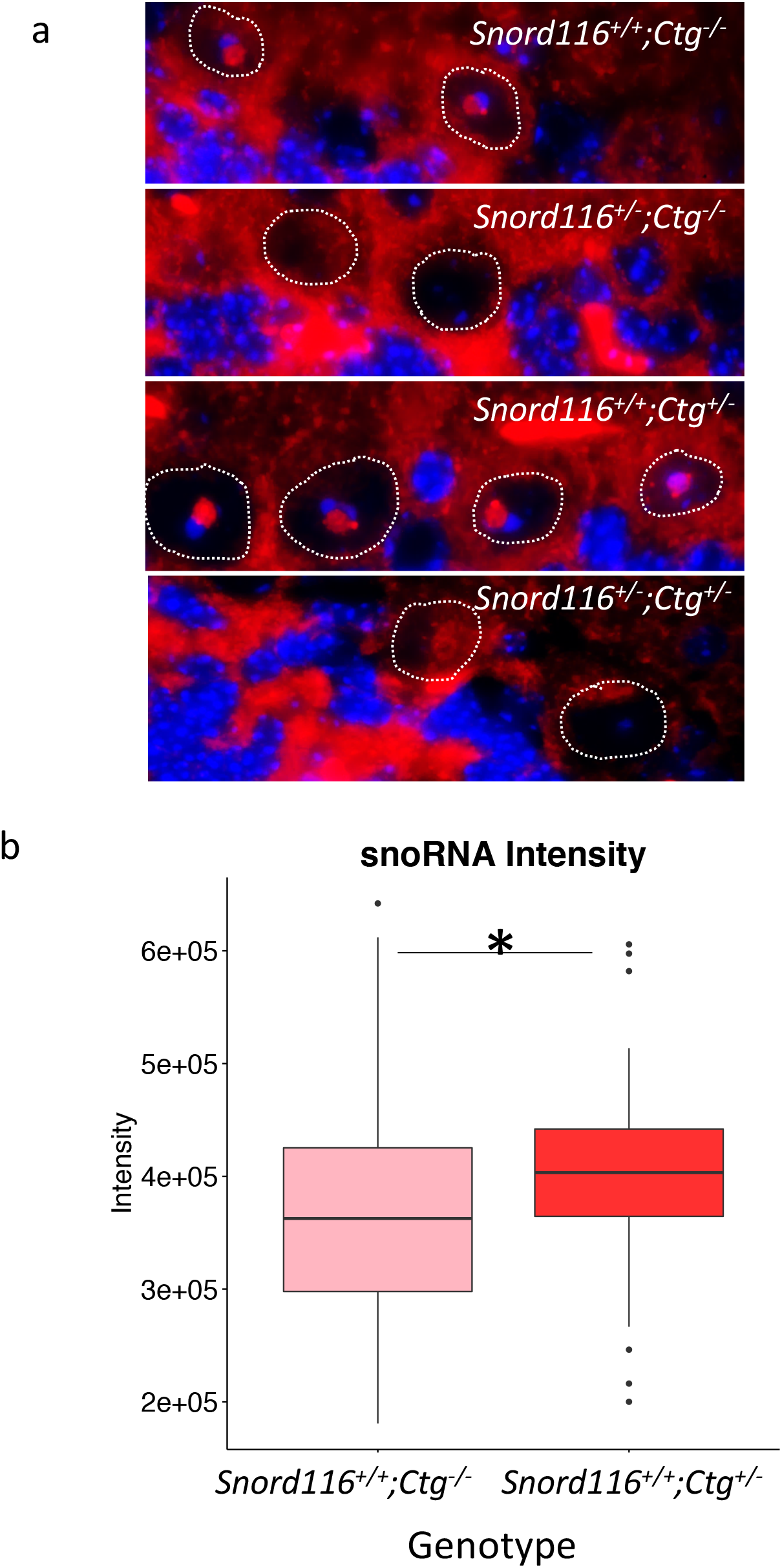
Transgenic *Snord116* colocalizes with endogenous *Snord116*, but does not localize to nucleoli in neurons lacking endogenous Snordllß expression. **(a)** RNA FISH for *Snord116* snoRNAs in Purkinje neurons, **(b)** Quantification of RNA FISH signal shows significantly stronger *Snord116* snoRNA signal in *Snord116^+/+^;Ctg^+/−^* Purkinje nucleoli compared to *Snord116^+/+^;Ctg^−/−^*. *p=0.006 by t-test.

### Neuron-specific splicing of transgenic *Snord116* requires *Rbfox3/NeuN*

The discrepancy between expression of the primary transcript and production of functional snoRNA products from the *Snord116* transgene led us to determine if splicing was the limiting factor in snoRNA processing. Using transgene-specific primers, we assessed splicing of transgenic *Snord116* in several neuronal and non-neuronal tissues of *Snord116^+/+m^,Ctg^+/−^* and *Snord116^+/−^;Ctg^+/−^* mice. RT-PCR analysis demonstrated that splicing of transgenic *Snord116* was restricted to neuronal tissues of both *Snord116^+/+^;Ctg^+/−^* and *Snord116^+/−^;Ctg^+/−^* mice (Fig. 4a) even though primary transcripts were detected in multiple tissue types (Fig. 2d). These results demonstrate that splicing may explain the tissue-specific differences between transcript expression and snoRNA localization, but not the deficiency of snoRNA processing in neurons of *Snord116^+/−^*;*;Ctg^+/−^* mice.

**Figure 4.**
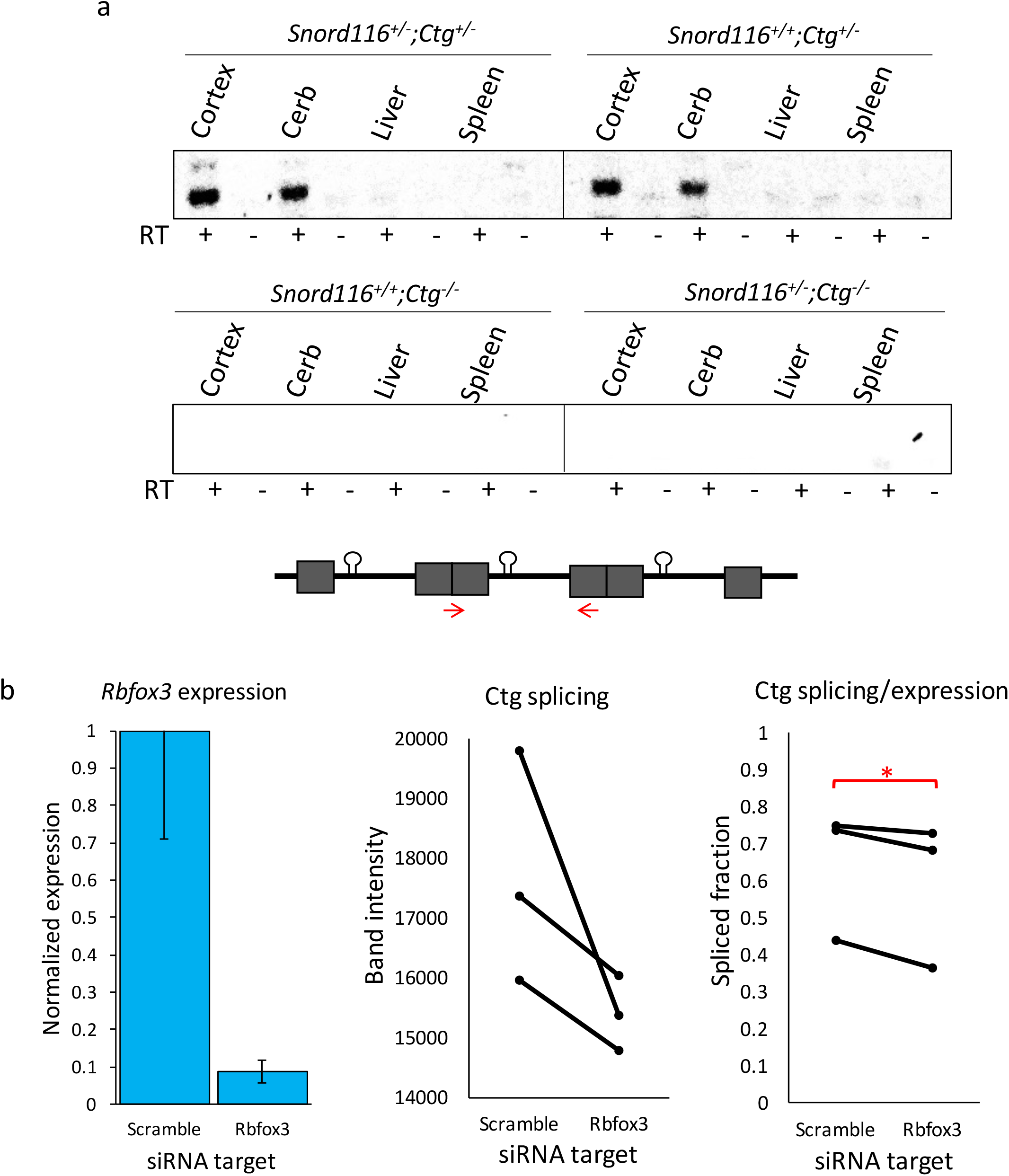
Splicing of transgenic *Snord116* is restricted to neuronal tissues and is reduced by Rbfox3 knockdown. **(a)** Splicing of transgenic *Snord116* is unique to neuronal tissues in both *Snord116^+/+^;Ctg^+/−^* and *Snord116^+/−^;Ctg^+/−^* mice, **(b)** Knockdown of *Rbfox3* expression in *Snord116^+/−^;Ctg^+/−^* NPC-derived neurons reduces splicing of the *Snord116* transgene. The spliced fraction of the expressed transgene is significantly lower with *Rbfox3* knockdown. *p=0.049 by paired t-test.

NeuN is commonly used in immunofluorescence staining as a neuron-specific marker, and has now been identified as the neuron-specific splicing factor, *Rbfox3* (23). To test the hypothesis that Rbfox3/NeuN regulates neuron-specific splicing of the *Snord116* transgene, we performed an siRNA knockdown of *Rbfox3* in neurons derived from *Snord116^+/−^;Ctg^+/−^* neural progenitor cells (NPCs), reducing *Rbfox3* expression to about 9% of levels detected in scramble siRNA treated NPCs (Figs. 4b and S3). The expression of the transgenic *Snord116* primary transcript was unaffected following *Rbfox3* siRNA knockdown, however the proportion of the primary transcript that was spliced (splicing/expression) was significantly reduced (Fig. 4b and S3). To further examine a role for Rbfox3 in *Snord116* splicing, we utilized a published dataset of RNA sequencing from *Rbfox3* knockout mouse cerebral cortex (24). Visualization of exon junctions within the *Snord116* locus by sashimi plot using the IGV browser demonstrated that loss of Rbfox3 leads to pronounced dysregulation of exon splicing between *Snord116* and *Ube3a* (Fig. S4). Together, these results demonstrate that *Rbfox3* levels affect neuronal *Snord116* splicing, and suggest that Rbfox3/NeuN is essential for the processing of intron-embedded snoRNAs.

### The transgenic host gene *116HG* RNA cloud localizes to the endogenous, but not the transgenic *Snord116* locus, only in wild-type neurons

In addition to the nucleolar snoRNAs, the spliced exons of the *Snord116* locus are retained as a RNA cloud that localizes to the site of transcription on the active paternal allele in wild-type neurons (22) (Fig. 1). We therefore asked whether the transgenically-encoded *116HG* localized to its own site of transcription using RNA FISH with a probe designed to detect the spliced junctions of both endogenous and transgenic *116HG.* Similarly to the *Snord116* snoRNA localization, *116HG* FISH signals were observed as a single nuclear cloud in both WT and *Snord116^+/+^;Ctg^+/−^* neurons, but not *Snord116^+/−^,Ctg^−/−^* or *Snord116^+/−^;Ctg^+/−^* neurons (Fig. 5a and Figs. S5 and S6). In mice of two different ages (5.5 months or 1 year), the single *116HG* RNA FISH signal was significantly larger in *Snord116^+/+^;Ctg^+/−^* compared to WT neurons (Fig. 5b), suggesting that the transgenic spliced *116HG* localized and contributed to the *116HG* RNA cloud on the endogenous paternal allele. By DNA FISH, distinct nuclear locations of the three alleles (endogenous maternal and paternal, plus transgene) were observed (Fig. 5c), demonstrating the absence of colocalization of the transgenic allele with the active paternal allele at the chromosomal level. Concordant with the lack of *Snord116* molecular rescue of the *116HG* RNA cloud and the *Snord116* snoRNAs, *Snord116+^/−^; Ctg+^-^* mice had a significantly lower body weight than *Snord116^+/+^;Ctg^−/−^* mice, similar to the weights observed in *Snord116^+/−^;Ctg^−/−^* mice (Fig. 6a) (14). Interestingly, the additive effect of endogenous and transgenic *Snord116* also resulted in decreased body weight compared to expression of the endogenous locus alone (WT), reaching significance at postnatal week 11, suggesting a potential dosage effect of *Snord116* on metabolism.

**Figure 5.**
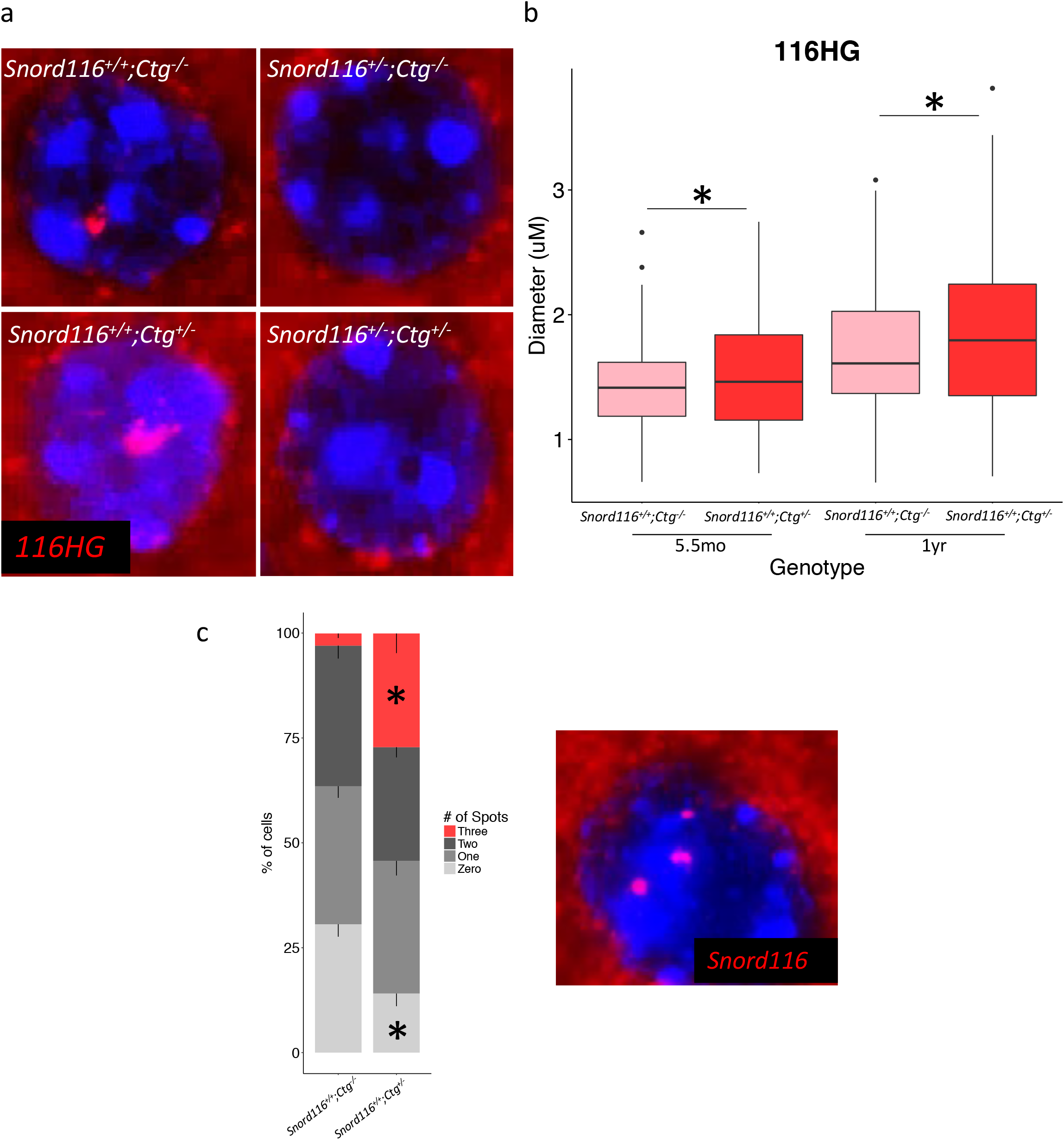
Transgenic *116HG* colocalizes with the endogenous *116HG* RNA cloud, but is not detected in the nuclei of neurons lacking endogenous *Snord116* expression. **(a)** RNA FISH for *116HG* in a cortical neuronal nucleus, **(b)** Quantification of RNA FISH signal shows a single significantly larger *116HG* RNA cloud in *Snord116^+/+^;Ctg^+/−^* neuronal nuclei compared to *Snord116^+/+^;Ctg^+/−^* T-test, p=0.002. **(c)** DNA FISH of *Snord116* in neuronal nuclei shows three detectable *Snordll6* loci in a significant proportion of *Snord116^+/+^;Ctg^+/−^* neuronal nuclei, indicating a lack of colocalization. *p<0.0001 by t-test.

**Figure 6.**
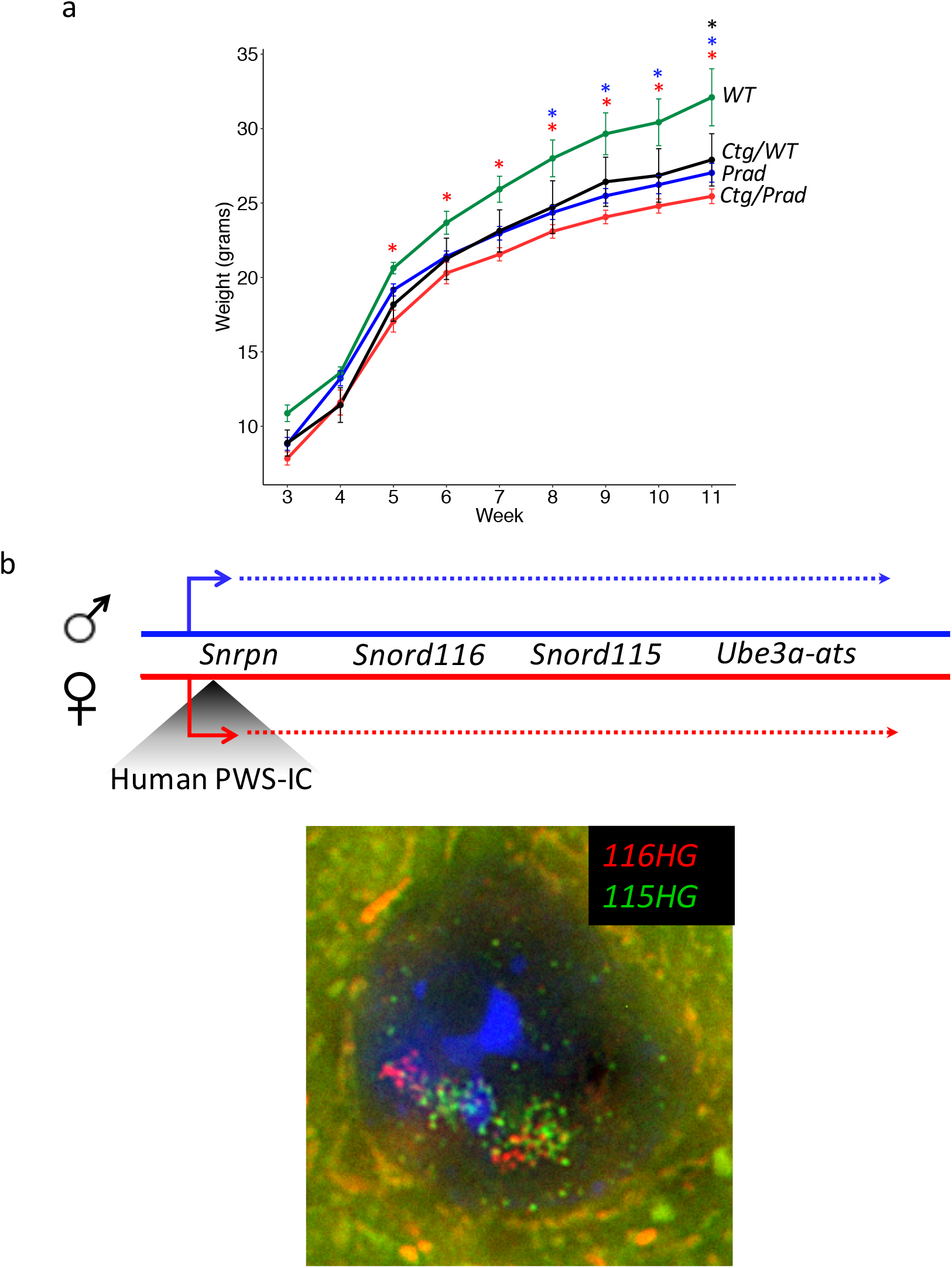
Transgenic *Snord116* does not rescue the weight phenotype observed in *Snord116^+/−^* mice. **(a)** Weight curves of all genotypes from *Snord116^+/−^;Ctg^−/−^* X *Snord116^+/+^;Ctg^+/−^* cross. Mice carrying the *Snord116* transgene, either on a WT or *Snord116^+/−^* background weigh significantly less than WT mice, and are phenotypically more similar to *Snord116^+/−^* mice. N = 4 *Snord116^+/+^Ctg^−/−^* (WT), 4 *Snord116^+/+^;Ctg^+/−^* (Ctg/WT), 13 *Snord116^+/−^;Ctg^−/−^* (Prad), 13 *Snord116^+/−^;Ctg^+/−^* (Ctg/Prad). *p<0.05 by repeated measures ANOVA, Benjamini-Hochberg post-hoc correction (results in Supplementary table 1). **(b)** Map of the 15qll-ql3 locus in PWS-IC^Hs^/+ mice. RNA FISH for *116HG* (red) and *115HG* (green) in a PWS-IC^Hs^/+ neuronal nucleus, with two decondensed alleles, shows the formation of two distinct RNA clouds for each cluster, indicating the requirement for a decondensed endogenous locus for the localization of the spliced *116HG.*

To test the hypothesis that a second requirement for *116HG* complementation is an active endogenous *Snord116* locus, we utilized another transgenic mouse model, in which the maternal mouse PWS-IC has been replaced by the human PWS-IC (PWS-IC^Hs^/+) (25). Because the human PWS-IC does not become imprinted in mice, the normally silent maternal allele undergoes chromatin decondensation and expresses *Snrpn*, snoRNAs and *Ube3a-ats* in neurons, similar to the paternal allele (Fig. 6b) (10).

In neuronal nuclei of PWS-IC^Hs^/+ mice, two distinct *116HG* RNA clouds were observed by RNA FISH, demonstrating that the requirement for *116HG* RNA localization is an active endogenous *Snord116* allele, not simply transcription of *Snord116.* Together, these results demonstrate that complementation by transgenic *Snord116* requires additional factors both *in trans (Rbfox3/NeuN)* and *in cis* (chromatin decondensation).

## Discussion

This study utilized a novel 27-copy *Snord116* transgenic mouse to characterize the requirements for *Snord116* processing and assess the molecular functional capacity of transgenic *Snord116* in the absence of endogenous paternal expression. We demonstrated that transgenic *Snord116* co-localizes with endogenous *Snord116* snoRNAs and *116HG* in the brain, however transgenic *Snord116* expression alone is not sufficient for the formation of functional RNAs from this locus, as they are not detected in tissues outside of the brain. Additionally, trafficking of transgenic RNAs is impaired in the absence of endogenous paternal *Snord116*, therefore nuclear RNA cloud formation is observed only in the presence of endogenous *Snord116* expression. Further, splicing of the *Snord116* transgene was restricted to the brain, despite expression in many tissues. This process is partially blocked by the knockdown of *Rbfox3* in NPC-derived neurons, suggesting a role for this neuron-specific splicing factor in the processing of *Snord116.*

Splicing represents a critical step in the regulation of gene expression in all tissues, however brain exhibits the highest levels of tissue specific alternative splicing (26–28). Such tissue-specific splicing reflects the intricate cellular connections and functional diversity within the brain and exemplifies the complex expression and regulatory networks observed in the brain and during development. Brain-specific splicing factors provide an important mechanism for the co-and post-transcriptional regulation involved in processes such as neurogenesis and synapse formation through the spatiotemporal control of RNA expression, processing, and localization (29,30). The Rbfox family of splicing factors are important in the regulation of development of the brain, with *Rbfox3* expression uniquely confined to mature postmitotic neurons (31). Neuronal co-expression and the presence of the (U)GCAUG binding motif (32,33) within the introns of the *Snord116* transcript support the role for Rbfox3 in the processing of *Snord116* in the brain. In addition, a recent study modeling the Rbfox3 regulatory network by crosslinking and immunoprecipitation followed by high-throughput sequencing (HITS-CLIP) identified significant peaks within the introns of *Ipw*, validating our Snord116-specific analysis of Rbfox3 activity (33).

Phenotypes associated with *Rbfox3* deficiency are relevant to neurodevelopment and sleep, and are likely the result of dysregulation of downstream targets (34,35). All three Rbfox splicing factors are downregulated in the brains of individuals with autism, suggesting a potential role for the dysregulation of this regulatory splicing network in the pathogenesis of autism and related neurodevelopmental disorders (33). Knockout of *Rbfox3* results in hippocampal synaptic deficits, and mutations of *Rbfox3* are implicated in neurodevelopmental delay (34). Interestingly, gene network analysis suggests a role for Rbfox3 in the release of neurotransmitters central to the circadian clock, and variants are associated with sleep latency (35). Paternal *Snord116* deficiency in PWS coincides with shorter nighttime sleep duration and daytime sleepiness, indicators of prolonged sleep latency (36,37). Electrophysiological examination of these sleep phenotypes revealed disrupted non-rapid eye movement (NREM) and rapid eye movement (REM) sleep in both *PWScr^m+/p-^* mice and PWS patients (36). In accordance with these phenotypic analyses, molecular characterization of *Snord116* demonstrated the diurnal regulation of the spliced *116HG*, which forms a significantly larger RNA cloud during light (sleep) hours, coinciding with downregulation of circadian genes, including *Per2, Per3, Bmal1*, and *Clock* (14). Furthermore, we recently identified >23,000 rhythmic methylated CpGs were observed in wild-type mouse cortex, of which 97% were lost or time-shifted in *Snord116^+/−^* littermates, including enhancers and promoters regulating genes with functions highly enriched for circadian entrainment and body weight (38).

The identification of *Snord116* as a splicing target of Rbfox3 supports this shared circadian phenotype as a downstream effect of altered *Snord116* processing.

*Ube3a*, another gene within the 15q11-q13 locus, is paternally imprinted in neurons, in which the paternal expression of the *Snrpn-Ube3a* transcript extends through the anti-sense to *Ube3a*, blocking transcription of paternal *Ube3a* and leading to maternal-specific expression (39–41). Interestingly, in Rbfox3/NeuN-negative cells of the suprachiasmatic nucleus (SCN), paternal *Ube3a* expression is observed (42). Due to its high level of GC-skew, transcription through *Snord116* results in the formation of R-loops, modulating the balance between chromatin state and transcription elongation. Stabilization of these R-loops by topotecan treatment stalls RNA polymerase II progression, blocking transcription through *Ube3a-ats* and resulting in biallelic *Ube3a* expression (22). Conversely, a study of genome stability demonstrated the ability of the ASF/SF2 splicing factor to interact with R-loops, repressing their formation through interaction with the nascent RNA (43). The Rbfox3-dependent silencing of paternal *Ube3a* in the SCN suggests that neuron-specific splicing of *Snord116* by Rbfox3 may be important for maintaining the balance of R-loop formation, promoting transcription through *Ube3a-ats* and paternal imprinting of *Ube3a* in neurons.

Lack of snoRNA and *116HG* formation in *Snord116^+/−^;Ctg^+/−^* neurons indicates a multi-level deficit in the production of functional RNAs from transgenic *Snord116.* Paternal *Snord116* is GC-skewed, resulting in R-loop formation, histone displacement, and chromatin decondensation upon neuronal transcription, and the specific localization of the *116HG* to its site of transcription at the decondensed paternal allele suggests a chromatin-state-dependent accumulation of *116HG* (22). The formation of two *116HG* RNA clouds in PWS-IC^Hs^/+ neuronal nuclei demonstrates the requirement for a decondensed endogenous allele for the localization of *Snord116*, suggesting that chromatin decondensation may mediate RNA-DNA interactions necessary to anchor the *116HG* to its proper subnuclear domain. Further study of the mechanisms of *Snord116* localization would benefit our understanding of the requirements for *Snord116* function and enable the development of more effective therapies in the future.

## Materials and Methods

### Mouse Husbandry

C57BL/6J (WT) and B6(Cg)-Snord116tm1.1Uta/J *(Snord116^+/−^)* mice were obtained from Jackson Labs (Bar Harbor, ME, USA). *Ctg^+/−^* mice were generated by the Mouse Biology Program (UC Davis). All mice were housed in a 12h light:12h dark, temperature controlled room and fed a standard diet of Picolab mouse chow 20 (PMI International, St Louis, MO, USA). Heterozygous deletion male mice *(Snord116^+/−^)* were bred with heterozygous *Ctg* transgenic females *(Ctg^+/−^)* to generate littermates of each of the following genotypes: *Snord116^+/+^Ctg^−/−^* (WT), *Snord116^+/−^;Ctg^−/−^* (Prad), *Snord116^+/+^;Ctg^+/−^* (Ctg/WT), *Snord116^+/−^;Ctg^+/−^* (Ctg/Prad). All mice used for this study were male and tissues were collected during light hours (ZT0-ZT12).

### RNA FISH and DNA FISH

*Snord116* BAC RP23-358G20 was ordered from BACPAC Resources (Children’s Hospital Oakland Research Institute). Nick translation of DIG labeled probes and DNA FISH were performed as described previously (10). RNA FISH was performed as described previously (10).

*116HG* probes = AATGCAACCCTTTTAACTCAG (Exiqon), pooled probes (Agilent). snoRNA probe = TTCCGATGAGAGTGGCGGTACAGA (Exiqon).

### Microscopy

Slides were imaged on an Axioplan 2 fluorescence microscope (Carl Zeiss) equipped with a Qimaging Retiga EXi high-speed uncooled digital camera and analyzed using iVision software (BioVision Technologies). Images were captured using a 100x oil objective and 1x camera zoom. *116HG* RNA cloud measurements were taken as two perpendicular cross-sections through the center of the RNA cloud and averaged for RNA cloud size. snoRNA intensity was measured as the sum of the intensity of each pixel divided by the area measured. All measurements were converted from pixel counts to microns according to objective and zoom (1px = 0.069μm). All measurements and spot counting were blinded to minimize bias.

### RT-PCR

RNA was isolated with the RNeasy mini kit (Qiagen) and cDNA was synthesized using the Quantitect reverse transcription kit (Qiagen). RT-PCR was performed using custom primers.

Ctg expression (transgene specific): F - taagcagagctggtttagtgaacc; R - aacagttcgatggagactcagttgg

Ctg splicing (transgene specific): F - cctgagttaaaaggcggccg; R - gccatttcctctgcatgttt Rbfox3 expression: F - aattttcccgaattgcccgaac; R - atgaagcagcacagacagacaa

### NPC and Neuron Culture

Embryos were dissected and cortices removed at E15. Neural progenitor cells were isolated as described previously (44) and cultured in NEP Complete media supplemented with Glutamax. To differentiate, neurospheres were dissociated with Accutase (Invitrogen) and filtered for a single cell suspension. Plates were coated with Poly-D-lysine (Sigma) and laminin (Invitrogen) and media was changed to Neurobasal with retinoic acid and BDNF.

### siRNA Knockdown

*Rbfox3* was knocked down using a pool of three Stealth RNAi siRNA or a negative control siRNA at 55 pmol RNAi per 60 mm dish (Life Technologies). Neurons were differentiated for 7 days followed by 3 days of knockdown. RNA was then extracted and evaluated for knockdown efficiency by qRT-PCR and expression/splicing by RT-PCR.

### Inverse PCR

Genomic DNA was isolated from a tail clipping using the Gentra Puregene kit (Qiagen). DNA was digested with DpnII and circularized by T4 ligation (Promega). Primers were designed to the known transgene sequence to amplify the unknown flanking genomic region. The PCR product was gel purified and sequenced by Sanger sequencing. Sequences flanking the Dpn II restriction site were mapped to the transgene and the genome flanking.

Inverse PCR primers: F - gatttccaagtctccaccccat; R – ggctatgaactaatgaccccgt

### RNA-seq Analysis

SRA files were downloaded from GEO (GSE84786) (24) and converted to fastq files using fastq-dump, splitting files for paired-end reads. Reads were trimmed for adapters and quality using the following parameters: ILLUMINACLIP:TruSeq3-PE.fa:2:30:10:8:T LEADING:15 TRAILING:15 SLIDINGWINDOW:4:15. An insert size of 200 bp was used based on Picard metrics and reads were aligned to mm10 using Tophat2. Bam index files were built using Picard Tools and stranded bed files were created and used to create bedgraph and bigwig files for visualization as custom tracks on the UCSC genome browser. Raw bam files were loaded into the IGV browser to create sashimi plots (MISO framework) (45,46).

## Acknowledgements

This work was supported by the NIH R01 NS076263, R56 NS076263-06 (JML), T32 GM007377 (RLc), IDDRC U54 HD079125, and the Foundation for Prader-Willi Research.

## Abbreviations

small nucleolar RNA (snoRNA); processed snoRNAs (psnoRNAs); Prader-Willi syndrome (PWS); Angelman syndrome (AS); imprinting center (iC); Prder-Willi critical region; (PWScr); non-coding RNAs (ncRNAs); uniparental disomy (UPD); microRNA (miRNA); ribosomal RNA (rRNA); fluorescence *in situ* hybridization (FISH); cytomegalovirus (CMV)

## References

1. Cassidy SB, Schwartz S, Miller JL, Driscoll DJ. Prader-Willi syndrome. Genet Med. 2012;14(1).

2. Runte M, Hüttenhofer A, Groß S, Kiefmann M, Horsthemke B, Buiting K. The IC-SNURF-SNRPN transcript serves as a host for multiple small nucleolar RNA species and as an antisense RNA for UBE3A. Hum Mol Genet. 2001;10(23):2687–700.

3. Landers M, Bancescu DL, Le Meur E, Rougeulle C, Glatt-Deeley H, Brannan C, et al. Regulation of the large (~1000 kb) imprinted murine Ube3a antisense transcript by alternative exons upstream of Snurf/Snrpn. Nucleic Acids Res. 2004;32(11):3480–92.

4. Sutcliffe JS, Nakao M, Christian S, Orstavik KH, Tommerup N, Ledbetter DH, et al. Deletions of a differentially methylated CpG island at the SNRPN gene define a putative imprinting control region. Nat Genet. 1994;8:52–8.

5. Buiting K, Saitoh S, Gross S, Dittrich B, Schwartz S, Nicholls RD, et al. Inherited microdeletions in the Angelman and Prader-Willi syndromes define an imprinting centre on human chromosome 15. Nat Genet. 1995;9:395–400.

6. Duker AL, Ballif BC, Bawle E V, Person RE, Mahadevan S, Alliman S, et al. Paternally inherited microdeletion at 15q 11.2 confirms a significant role for the SNORD116 C/D box snoRNA cluster in Prader-Willi syndrome. Eur J Hum Genet. 2010;18(10):1196–201.

7. de Smith AJ, Purmann C, Walters RG, Ellis RJ, Holder SE, Van Haelst MM, et al. A deletion of the HBII-85 class of small nucleolar RNAs (snoRNAs) is associated with hyperphagia, obesity and hypogonadism. Hum Mol Genet [Internet]. 2009;18(17):3257–65. Available from: http://hmg.oxfordjournals.org/

8. Sahoo T, Del Gaudio D, German JR, Shinawi M, Peters SU, Person RE, et al. Prader-Willi phenotype caused by paternal deficiency for the HBII-85 C/D box small nucleolar RnA cluster. Nat Genet. 2008;40(6):719–21.

9. Bieth E, Eddiry S, Gaston V, Lorenzini F, Buffet A, Auriol FC, et al. Highly restricted deletion of the SNORD116 region is implicated in Prader-Willi Syndrome. Eur J Hum Genet. 2014;(10):2014–103.

10. Leung KN, Vallero RO, Dubose AJ, Resnick JL, Lasalle JM. Imprinting regulates mammalian snoRNA-encoding chromatin decondensation and neuronal nucleolar size. Hum Mol Genet. 2009;18(22):4227–38.

11. Vitali P, Royo H, Marty V, Bortolin-Cavaille M-L, Cavaille J. Long nuclear-retained non-coding RNAs and allele-specific higher-order chromatin organization at imprinted snoRNA gene arrays. J Cell Sci [Internet]. 2010;123(1):70–83. Available from: http://jcs.biologists.org/cgi/doi/10.1242/jcs.054957

12. Cavaille J, Buiting K, Kiefmann M, Lalande M, Brannan CI, Horsthemke B, et al. Identification of brain-specific and imprinted small nucleolar RNA genes exhibiting an unusual genomic organization. Proc Natl Acad Sci U S A [Internet]. 2000;97(26):14311–6. Available from: http://www.pubmedcentral.nih.gov/articlerender.fcgi?artid=18915&tool=pmcentrez&rendertype=abstract

13. Falaleeva M, Stamm S. Processing of snoRNAs as a new source of regulatory non-coding RNAs: SnoRNA fragments form a new class of functional RNAs. BioEssays. 2013;

14. Powell WT, Coulson RL, Crary FK, Wong SS, Ach RA, Tsang P, et al. A Prader-Willi locus lncRNA cloud modulates diurnal genes and energy expenditure. Hum Mol Genet. 2013;22(21).

15. Bervini S, Herzog H. Mouse models of Prader-Willi Syndrome: A systematic review. Frontiers in Neuroendocrinology. 2013.

16. Purtell L, Qi Y, Campbell L, Sainsbury A, Herzog H. Adult-onset deletion of the Prader-Willi syndrome susceptibility gene Snord116 in mice results in reduced feeding and increased fat mass. Transl Pediatr. 2017;6(2):88–97.

17. Polex-Wolf J, Lam BYH, Larder R, Tadross J, Rimmington D, Bosch F, et al. Hypothalamic loss of Snord116 recapitulates the hyperphagia of Prader-Willi syndrome. J Clin Invest. 2018;128(3):960–9.

18. Skryabin B V., Gubar L V., Seeger B, Pfeiffer J, Handel S, Robeck T, et al. Deletion of the MBII-85 snoRNA gene cluster in mice results in postnatal growth retardation. PLoS Genet. 2007;3(12).

19. Ding F, Li HH, Zhang S, Solomon NM, Camper SA, Cohen P, et al. SnoRNA Snord116 (Pwcr1/MBll-85) deletion causes growth deficiency and hyperphagia in mice. PLoS One. 2008;

20. Rozhdestvensky TS, Robeck T, Galiveti CR, Raabe CA. Maternal transcription of non-protein coding RNAs from the PWS-critical region rescues growth retardation in mice. Nature [Internet]. 2016;6. Available from: http://dx.doi.org/10.1038/srep20398

21. Tsai TF, Jiang YH, Bressler J, Armstrong D, Beaudet AL. Paternal deletion from Snrpn to Ube3a in the mouse causes hypotonia, growth retardation and partial lethality and provides evidence for a gene contributing to Prader-Willi syndrome. Hum Mol Genet. 1999;

22. Powell WT, Coulson RL, Gonzales ML, Crary FK, Wong SS, Adams S, et al. R-loop formation at Snord116 mediates topotecan inhibition of Ube3a-antisense and allele-specific chromatin decondensation. Proc Natl Acad Sci U S A. 2013;110(34).

23. Kim KK, Adelstein RS, Kawamoto S. Identification of Neuronal Nuclei (NeuN) as Fox-3, a New Member of the Fox-1 Gene Family of Splicing Factors. J Biol Chem. 2009;284(45):31052–61.

24. Wang H-Y, Hsieh P-F, Huang D-F, Chin P-S, Chou C-H, Tung C-C, et al. RBFOX3/NeuN is Required for Hippocampal Circuit Balance and Function. Nat Publ Gr. 2015;5(17383).

25. Johnstone KA, DuBose AJ, Futtner CR, Elmore MD, Brannan CI, Resnick JL. A human imprinting centre demonstrates conserved acquisition but diverged maintenance of imprinting in a mouse model for Angelman syndrome imprinting defects. Hum Mol Genet. 2006;

26. de la Grange P, Gratadou L, Delord M, Dutertre M, Auboeuf D. Splicing factor and exon profiling across human tissues. Nucleic Acids Res. 2010;38(9):2825–38.

27. Xu Q, Modrek B, Lee C. Genome-wide detection of tissue-specific alternative splicing in the human transcriptome. Nucleic Acids Res. 2002;30(17):3754–66.

28. Yeo G, Holste D, Kreiman G, Burge CB. Variation in alternative splicing across human tissues. Genome Biol. 2004;5(10).

29. Zheng S, Black DL. Alternative Pre-mRNA Splicing in Neurons, Growing Up and Extending Its Reach. Trends Genet. 2014;29(8):442–8.

30. Chih B, Gollan L, Scheiffele P. Alternative Splicing Controls Selective Trans-Synaptic Interactions of the Neuroligin-Neurexin Complex. Neuron. 2006;51:171–8.

31. Duan W, Zhang Y-P, Hou Z, Huang C, Zhu H, Zhang C-Q, et al. Novel Insights into NeuN: from Neuronal Marker to Splicing Regulator. Mol Neurobiol. 2015;

32. Minovitsky S, Gee SL, Schokrpur S, Dubchak I, Conboy JG. The splicing regulatory element, UGCAUG, is phylogenetically and spatially conserved in introns that flank tissue-specific alternative exons. Nucleic Acids Res. 2005;33(2):714–24.

33. Weyn-vanhentenryck SM, Mele A, Yan Q, Sun S, Farny N, Zhang Z, et al. HITSCLIP and integrative modeling define the Rbfox splicing-regulatory network linked to brain development and autism. Cell Rep. 2014;6(6):1139–52.

34. Utami KH, Hillmer AM, Aksoy I, Chew EG, Teo AS, Zhang Z, et al. Detection of chromosomal breakpoints in patients with developmental delay and speech disorders. PLoS One [Internet]. 2014;9(6):e90852. Available from: https://www.ncbi.nlm.nih.gov/pubmed/24603971

35. Amin N, Allebrandt K V., van der Spek A, Muller-Myhsok B, Hek K, Teder-Laving M. Genetic variants in RBFOX3 are associated with sleep latency. Eur J Hum Genet. 2016;

36. Lassi G, Priano L, Maggi S, Garcia-Garcia C, Balzani E, El-Assawy N, et al. Deletion of the Snord116/SNORD116 Alters Sleep in Mice and Patients with Prader-Willi Syndrome. Sleep [Internet]. 2016;39(3):637–44. Available from: https://academic.oup.com/sleep/article-lookup/doi/10.5665/sleep.5542

37. Gibbs S, Wiltshire E, Elder D. Nocturnal sleep measured by actigraphy in children with Prader-Willi syndrome. J Pediatr [Internet]. 2013;162(4):765–9. Available from: http://dx.doi.org/10.1016/j.jpeds.2012.09.019

38. Coulson RL, Yasui DH, Dunaway K, Laufer BI, Vogel Ciernia A, Zhu Y, et al. Snord116-dependent diurnal rhythm of DNA methylation in mouse cortex. Nat Commun. In Press 2018; 10.1038/s41467-018-03676-0

39. Rougeulle C, Cardoso C, Fontes M, Colleaux L, Lalande M. An imprinted antisense RNA overlaps UBE3A and a second maternally expressed transcript. Nat Genet. 1998;19:15–6.

40. Meng L, Person RE, Beaudet AL. Ube3a-ATS is an atypical RNA polymerase II transcript that represses the paternal expression of Ube3a. Hum Mol Genet. 2012;21(13):3001–12.

41. Meng L, Person RE, Huang W, Zhu PJ, Costa-Mattioli M, Beaudet AL. Truncation of Ube3a-ATS Unsilences Paternal Ube3a and Ameliorates Behavioral Defects in the Angelman Syndrome Mouse Model. PLoS Genet. 2013;9(12).

42. Jones KA, Han JE, DeBruyne JP, Philpot BD. Persistent neuronal Ube3a expression in the suprachiasmatic nucleus of Angelman syndrome model mice. Sci Rep [Internet]. 2016;6(1):28238. Available from: http://www.nature.com/articles/srep28238

43. Aguilera A, Garcia-Muse T. R loops: from transcription byproducts to threats to genome stability. Mol Cell [Internet]. 2012/05/01. 2012;46(2):115–24. Available from: http://www.ncbi.nlm.nih.gov/pubmed/22541554

44. Hutton SR, Pevny LH. Isolation, culture, and differentiation of progenitor cells from the central nervous system. Cold Spring Harb Protoc. 2008;3(11).

45. Katz Y, Wang ET, Airoldi EM, Burge CB. Analysis and design of RNA sequencing experiments for identifying isoform regulation. Nat Methods. 2010;7(12):1009–15.

46. Katz Y, Wang ET, Silterra J, Schwartz S, Wong B, Thorvaldsdóttir H, et al. Quantitative visualization of alternative exon expression from RNA-seq data. Bioinformatics. 2015;31 (14):2400–2.

